# No impact of functional connectivity of the motor system on the resting motor threshold: A replication study

**DOI:** 10.1101/2020.10.26.354886

**Authors:** Melina Engelhardt, Darko Komnenić, Fabia Roth, Leona Kawelke, Carsten Finke, Thomas Picht

**Author notes:** **Correspondence:** Melina Engelhardt.

## Abstract

The physiological mechanisms of corticospinal excitability and factors influencing its measurement with transcranial magnetic stimulation are still poorly understood. A recent study reported an impact of functional connectivity between the primary motor cortex and dorsal premotor cortex on the resting motor threshold of the dominant hemisphere. We aimed to replicate these findings in a larger sample of 38 healthy right-handed subjects with data from both hemispheres. Resting-state functional connectivity was assessed between the primary motor cortex and five a-priori defined motor-relevant regions on each hemisphere as well as interhemispherically between both primary motor cortices. Following the procedure by the original authors, we included age, the cortical grey matter volume and coil to cortex distance as further predictors in the analysis. We report replication models for the dominant hemisphere as well as an extension to data from both hemispheres and support the results with Bayes factors. Functional connectivity between the primary motor cortex and dorsal premotor cortex did not explain variability in the resting motor threshold and we obtained moderate evidence for the absence of this effect. In contrast, coil to cortex distance could be confirmed as an important predictor with strong evidence. These findings contradict the previously proposed effect, thus questioning the notion of the dorsal premotor cortex playing a major role in modifying corticospinal excitability.

## 1 Introduction

Resting-motor threshold (RMT) is a fundamental measurement in transcranial magnetic stimulation (TMS) studies. It is commonly used as an indicator of cortical excitability and as a basic dosing unit for TMS-based therapeutic interventions. These interventions have seen usage in multiple disciplines ranging from studies in motor cortical mapping, depression, language and vision (for an overview of different stimulation protocols see Lefaucheur et al. (2014)). Despite its prevalent use, RMT’s underlying physiological mechanisms and modulating factors are still poorly understood (Herbsman et al. 2009; Hübers et al. 2012; Wassermann 2002). To assure an accurate RMT assessments, specifically when used as an outcome measurement to assess treatment effect, potential confounders need to be identified and their influence minimized.

The RMT is defined as the smallest stimulation intensity to reliably elicit motor evoked potentials in a target muscle using TMS (Caramia et al. 1989; P. Rossini, Barker, and Berardelli 1994; P. M. Rossini et al. 2015; Rothwell, J. C., Hallett, M., Berardelli, A., Eisen, A., Rossini, P., & Paulus 1999). It is used to capture excitability of stimulated cortical motor areas. Specifically, it reflects transsynaptic activation of corticospinal neurons as it can be modulated by changing conductivity of presynaptic sodium or calcium channels (Ziemann et al. 1996).

Several studies (Bhandari et al. 2016; Latorre et al. 2019; Wassermann 2002) have shown a substantial variability in RMT between and within healthy subjects. While the impact of methodological factors such as the TMS equipment, use of neuronavigation software and algorithms used to assess RMT is well established, the effects of structural and functional factors are still poorly understood (Herbsman et al. 2009; Hübers et al. 2012; Rosso et al. 2017). Recent studies have shown a positive correlation of RMT with subject age after maturation of the white matter, a relationship potentially mediated by a reduction of cortical volume and increase in coil-cortex distance (CCD; Bhandari et al. 2016; Rosso et al. 2017). Independent of age, CCD has been replicated as an important predictor of the RMT (McConnell et al. 2001; Kozel et al. 2000; Stokes et al. 2005). Further, cortical thickness of the motor hand knob was positively correlated with RMT in one study (List et al. 2013). Results are conflicting regarding the impact of white matter properties assessed using diffusion tensor imaging, e.g. fractional anisotropy (FA). Initial results (Klöppel et al. 2008) showing an inverse relationship between RMT and FA could not be replicated in subsequent studies (Herbsman et al. 2009; Hübers et al. 2012).

Rosso et al. (2017) were the first to study the impact of functional connectivity (FC) measured with resting-state functional magnetic resonance imaging (rsfMRI) on RMT, thereby including a measure of functional integration of motor information. They predicted RMTs of the dominant hemisphere with FC between the primary motor cortex (M1) and supplementary motor area (SMA), pre-SMA, dorsal premotor cortex (PMd), primary somatosensory cortex (S1) and the contralateral M1 using data of 21 participants. The impact of FC was then compared against known predictors such as age and CCD, as well as other factors such as FA and the cortical volume of these regions. The analysis showed a negative correlation between FC M1-PMd and RMT, which was confirmed in a multiple regression analysis including age, CCD and the cortical volume of the dominant hemisphere as well. The authors therefore concluded that cortical excitability of M1 is critically impacted by integration of information from PMd via cortico-cortical connections.

The aim of this study was to replicate these findings on the impact of FC M1-PMd in a larger sample and to assess their validity for the non-dominant hemisphere. We matched our sample in terms of age and gender distribution and followed the experimental design outlined by Rosso et al. (2017). We deviated from their paradigm only by using an atlas for delineation of the seed regions and focusing on the FC analysis, thus not investigating the impact of FA. Rosso et al. (2017) were contacted to inquire about details of the fMRI preprocessing and experimental setting, but were not included in any other way in this study. After this initial contact, we further included an exploratory analysis of the impact of the timing between the MRI and TMS procedure on our results.

## 2 Materials and Methods

As the present study was a replication attempt, we followed the experimental and analysis procedures of Rosso et al. (2017) as closely as possible. The software and protocols used for acquisition of the MRI data were similar to those used in Rosso et al. (2017) and analysis was identical. Remaining differences are specifically stated as such in the following methods. One deviation that became apparent only after contacting Rosso et al. (2017) was differences in the timing of the MRI and TMS procedures. While MRI and TMS procedures were performed consecutively in the study by Rosso et al. (2017), only a subset of our sample received both measures on the same day. We tried to account for these differences by including an exploratory analysis of this subset.

### 2.1 Participants

Thirty-eight healthy, right-handed subjects (age mean ± SD: 37.5 ± 13.8 years, 21 females) participated in the study. Seven of these subjects (age mean ± SD: 41.9 ± 18.5 years, 5 females) received the MRI immediately before the TMS procedure. Handedness was assessed with the Edinburgh Handedness Inventory (Oldfield 1971). Data was derived from two parallel studies (EA4/015/18, EA4/070/17) conducted at Charité. The inclusion criteria were (i) no history of neurological or psychiatric illness, (ii) age older than 18 years, (iii) no contraindications for TMS or MRI assessment, (iv) ability to provide written informed consent, (v) right-handedness. All study procedures were approved by the local ethics committee and the study was conducted in accordance with the Declaration of Helsinki. All subjects provided their written informed consent.

### 2.2 MRI

#### 2.2.1 Image Acquisition

MRI scans were performed on a Siemens 3-T Magnetom Trio MRI scanner (Siemens AG, Erlangen, Germany) with a 32-channel head coil. The MRI protocol took approximately 20 minutes and comprised a T1-weighted anatomical MPRAGE sequence (TR = 2530 ms; TE = 4.94 ms; TI = 1100 ms; flip angle = 7°; voxel size = 1 x 1 x 1 mm; 176 slices) and a resting-state fMRI sequence (TR = 2000 ms; TE = 30 ms; flip angle = 78°; voxel size = 3 x 3 x 3 mm; 238 volumes). For the rsfMRI sequence, subjects were instructed to close their eyes and let their thoughts flow freely.

#### 2.2.2 Rs-fMRI functional connectivity

Analysis of the rsfMRI functional connectivity was performed using the SPM-based Toolbox CONN (Version 18b; Whitfield-Gabrieli and Nieto-Castanon 2012). The functional and structural images were pre-processed using CONNs default preprocessing pipeline (Nieto-Castanon 2020). This includes the following steps: Functional images were realigned to the first scan of the sequence and then slice-time corrected. Potential outlier scans with framewise displacement above 0.5 mm or global BOLD signal changes above 3 standard deviations (according to the “conservative” standard in CONN) were identified. Anatomical and functional images were then normalized into MNI space and segmented into grey matter, white matter and cerebrospinal fluid. Finally, functional data were smoothed using a Gaussian kernel of 8mm full width half maximum. The default denoising pipeline as implemented in CONN (Nieto-Castanon 2020) was used subsequently. The performed procedures consist of a regression to remove potentially confounding components from white matter or cerebrospinal fluid, subject motion and previously identified outlier scans to improve the signal-to-noise ratio. The data were then band-pass filtered to retain frequencies from 0.008 to 0.1 Hz.

Following preprocessing, ROI-to-ROI functional connectivity matrices were computed by selecting the corresponding option within the first-level analysis segment in the CONN toolbox. Each element of the connectivity matrices represents a Fisher’s z-transformed bivariate correlation between a pair of ROI BOLD timeseries for one subject (Nieto-Castanon 2020). Deviating from Rosso et al. (2017), the Human Motor Area Template (Mayka et al. 2006) was used to define the ROIs included in the analysis in MNI space. This approach was chosen as it presents an objective, but time-efficient way to delineate ROIs in a larger number of subjects. Further, we decided to use this specific atlas as it matches the regions included in the original article with the inclusion of one additional ROI in the ventral premotor cortex (PMv). The following ROI-to-ROI functional connectivity values were included in the analysis within each hemisphere: M1-S1, M1-SMA, M1-preSMA, M1-PMd, M1-PMv. Additionally, interhemispheric functional connectivity was measured between right M1 and left M1 (M1-M1).

#### 2.2.3 Cortical gray matter volume

The cortical grey matter volume of each hemisphere was analyzed with Freesurfer (Version 7.1.0, http://surfer.nmr.mgh.harvard.edu/) using the recon-all command. Briefly, this procedure includes motion correction, removal of non-brain tissue, Talairach transformation, segmentation of grey and white matter structures, intensity normalization and cortical parcellation (Reuter et al. 2012; Fischl and Dale 2000; Fischl 2004).

#### 2.2.4 Coil-to-cortex distance

For measurement of the CCD, individual structural MRIs were analyzed using itk-SNAP (Version 3.8.0, www.itksnap.org; Yushkevich et al. 2006). The hand knob was localized for each hemisphere on the brain surface and the shortest distance between the cortical surface of the hand knob and the surface of the scalp was assessed.

### 2.3 Neuronavigated TMS

NTMS was applied using a Nexstim NBS5 stimulator (Nexstim, Helsinki, Finland) with a figure-of-eight coil (outer diameter: 70mm). Each subject’s structural MRI was used as a subject-specific navigational dataset. Motor evoked potentials were recorded in a belly-tendon fashion from the first dorsal interosseous muscles of both hands with disposable Ag/AgCl surface electrodes (Neuroline 700; Ambu, Ballerup, Denmark). The ground electrode was attached to the left palmar wrist. Subjects were instructed to sit comfortably in the chair and relax their hand muscles. Muscle activity was monitored to assure relaxation of the muscle, with a maximum tolerated baseline activity of 10 μV. The stimulation site, electric field direction and angulation consistently eliciting the largest motor evoked potentials in the target muscle was defined as the hotspot for stimulation and stored in the system. For this point, RMT was defined according to the Rossini-Rothwell method (Rossini, Barker, and Berardelli 1994; Rothwell et al. 1999) as lowest stimulation intensity to elicit motor evoked potentials larger than 50 μV in at least 5 out of 10 trials. RMT was recorded as a percentage of the maximum stimulator output.

### 2.4 Statistical Analysis

Statistical analyses were conducted in R Studio (Version 1.3.1073, http://www.rstudio.com/). Analysis was divided to first replicate results for the dominant hemisphere only (replication analysis) and second, to extend these findings to the whole dataset with data from both hemispheres (extended analysis). Finally, we tested the multiple regression model for the dominant hemisphere and linear mixed model for both hemispheres for the subset of participants (n = 7) that received the TMS procedure directly after the MRI. These last analyses should be interpreted with caution due to the small sample size of this subset of the data. Yet, we decided to include these illustrative analyses to give some idea about the impact of the timing between MRI and TMS as procedural deviation between both studies.

To assess the relationship between RMT and all included predictors alone, we replicated the correlation analyses of Rosso et al. (2017) for the data of the dominant hemisphere. Correlation coefficients, 95%-confidence intervals (CIs) and p-values are stated in Table 1. For the extended analysis, these relationships were quantified by linear mixed models with subjects as random intercepts. Estimates for fixed effects with 95%-CIs are presented together with t- and p-values approximated with Satterthwaite’s method (Table 2).

**Table 1.** Correlation coefficients for the dominant hemisphere.

| Dependent variable | Independent variable | Correlation coefficient<sup>a</sup> | T value | P value | BF<sub>10</sub> |
| --- | --- | --- | --- | --- | --- |
| RMT | CCD | 0.626<br>[0.383, 0.788] | 4.813 | < 0.001* | 784.65 |
|  | Age | 0.066<br>[-0.260, 0.377] | 0.394 | 0.696 | 0.39 |
|  | Grey matter volume | -0.187<br>[-0.478, 0.141] | -1.144 | 0.260 | 0.63 |
|  | FC M1-M1 | -0.130<br>[-0.432, 0.198] | -0.787 | 0.436 | 0.47 |
|  | FC M1-S1 | 0.043<br>[-0.281, 0.358] | 0.257 | 0.799 | 0.37 |
|  | FC M1-SMA | -0.156<br>[-0.453, 0.172] | -0.950 | 0.348 | 0.53 |
|  | FC M1-preSMA | 0.019<br>[-0.303, 0.336] | 0.112 | 0.911 | 0.36 |
|  | FC M1-PMd | 0.041<br>[-0.282, 0.356] | 0.249 | 0.805 | 0.37 |
|  | FC M1-PMv | 0.104<br>[-0.223, 0.410] | 0.627 | 0.535 | 0.43 |
| Age | Grey matter volume | -0.557<br>[-0.744, -0.289] | -4.027 | < 0.001* | 114.69 |

<sup>a</sup>Presented with 95% confidence intervals.
\*P-values below 0.05 were considered significant.

**Table 2.**
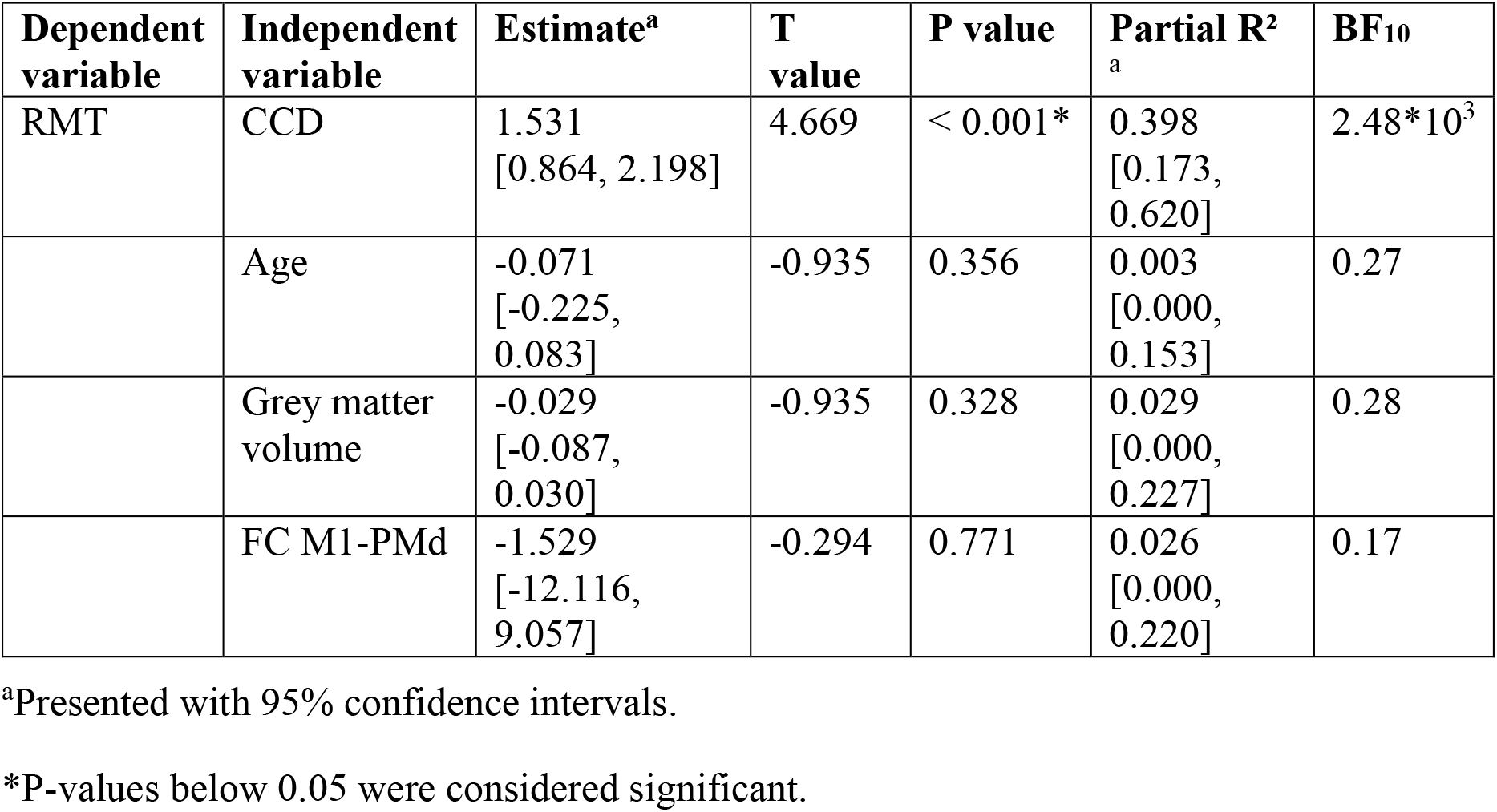
Multiple regression model for the dominant hemisphere.

| Dependent variable | Independent variable | Estimate<sup>a</sup> | T value | P value | Partial R<sup>2</sup><sub>a</sub> | BF<sub>10</sub> |
| --- | --- | --- | --- | --- | --- | --- |
| RMT | CCD | 1.531<br>[0.864, 2.198] | 4.669 | < 0.001* | 0.398<br>[0.173, 0.620] | 2.48*10<sup>3</sup> |
|  | Age | -0.071<br>[-0.225, 0.083] | -0.935 | 0.356 | 0.003<br>[0.000, 0.153] | 0.27 |
|  | Grey matter volume | -0.029<br>[-0.087, 0.030] | -0.935 | 0.328 | 0.029<br>[0.000, 0.227] | 0.28 |
|  | FC M1-PMd | -1.529<br>[-12.116, 9.057] | -0.294 | 0.771 | 0.026<br>[0.000, 0.220] | 0.17 |

<sup>a</sup>Presented with 95% confidence intervals.
\*P-values below 0.05 were considered significant.

In the replication analysis, we calculated the multiple linear regression model of Rosso et al. (2017) with RMT as dependent variable and age, CCD, the cortical volume of the hemisphere and FC M1-PMd as independent variables (Table 3). Estimates for regression coefficients with 95%-CIs are given together with t and p-values. Additionally, we computed the variance explained by the model R^2^ as well as partial R^2^ for each predictor with their respective 95%-CIs. In the extension analysis, we calculated a linear mixed model with RMT as dependent variable and age, CCD, the cortical grey matter volume of the hemisphere, hemisphere (0 = dominant, 1 = non-dominant) and FC M1-PMd as fixed effects (Table 4). Subjects were included as random effect. Estimates for fixed effects with 95%-CIs are given together with t- and p-values approximated with Satterthwaite’s method. Further, R^2^(Model) and partial R^2^ for each fixed effect with the respective 95%-CIs were computed.

**Table 3.** Linear mixed models with single variables using data from both hemispheres.

| **Dependent variable** | **Independent variable** | **Estimate<sup>a</sup>** | **T value<sup>b</sup>** | **P value<sup>b</sup>** | **BF<sub>10</sub>** |
| --- | --- | --- | --- | --- | --- |
| RMT | CCD | 1.448<br>[0.912, 1.976] | 5.452 | 0.001* | 1.59*10<sup>4</sup> |
|  | Age | 0.026<br>[-0.111, 0.162] | 0.378 | 0.708 | 0.12 |
|  | Grey matter volume | -0.022<br>[-0.080, 0.037] | -0.771 | 0.445 | 0.15 |
|  | FC M1-M1 | -4.039<br>[-11.157, 3.079] | -1.141 | 0.261 | 0.22 |
|  | FC M1-S1 | 2.014<br>[-1.469, 5.491] | 1.152 | 0.254 | 0.22 |
|  | FC M1-SMA | 1.910<br>[-3.727, 7.368] | 0.699 | 0.487 | 0.15 |
|  | FC M1-preSMA | -0.043<br>[-6.421, 6.329] | -0.013 | 0.989 | 0.12 |
|  | FC M1-PMd | -0.047<br>[-5.445, 5.256] | -0.017 | 0.986 | 0.12 |
|  | FC M1-PMv | -0.429<br>[-6.762, 6.041] | -0.137 | 0.891 | 0.12 |
|  | Hemisphere | -1.079<br>[-2.358, 0.201] | -1.695 | 0.098 | 0.46 |
| Grey matter volume | Age | -1.2847<br>[-1.907, -0.663] | -4.153 | < 0.001* | 140.31 |

<sup>a</sup>Presented with 95% confidence intervals.
<sup>b</sup>T and p values were approximated with Satterthwaite's method.
\*P-values below 0.05 were considered significant.

**Table 4.** Combined linear mixed model for both hemispheres.

| Dependent variable | Independent variable | Estimate<sup>a</sup> | T value<sup>b</sup> | P value<sup>b</sup> | Partial R<sup>2</sup><sub>a</sub> | BF<sub>10</sub> |
| --- | --- | --- | --- | --- | --- | --- |
| RMT | CCD | 1.468<br>[0.928, 1.999] | 5.508 | < 0.001* | 0.425<br>[0.271, 0.574] | 1.8*10<sup>4</sup> |
|  | Age | 0.034<br>[0.000, 0.156] | -1.080 | 0.287 | -0.070<br>[-0.198, 0.061] | 0.2 |
|  | Grey matter volume | 0.021<br>[0.000, 0.131] | -0.823 | 0.415 | -0.022<br>[-0.076, 0.032] | 0.16 |
|  | FC M1-PMd | 0.810<br>[-4.087, 5.710] | 0.329 | 0.743 | 0.001<br>[0.000, 0.074] | 0.12 |
|  | Hemisphere | -0.994<br>[-2.214, 0.227] | -1.638 | 0.110 | 0.016<br>[0.000, 0.118] | 0.42 |

<sup>a</sup>Presented with 95% confidence intervals.
<sup>b</sup>T and p values were approximated with Satterthwaite's method.
\*P-values below 0.05 were considered significant.

To assure interpretability of the results of regression and mixed models, we calculated variance inflation factors as a measure of collinearity between predictors in each model. A variance inflation factor < 5 suggests no collinearity between predictors. All models met this criterium. As in the original study, p-values ≤ 0.05 were considered significant.

While using these analyses with null hypothesis significance testing allows comparison with Rosso et al. (2017), it does not allow for rejection of the alternative hypothesis (Dienes 2011; 2014). However, judgement of evidence for or against the null hypothesis is crucial to decide whether a replication was successful. To quantify this evidence, we calculated Bayes factors (BF_10_) expressing evidence for the alternative hypothesis relative to the null hypothesis given the data. Thus, a Bayes factor > 1 provides anecdotal evidence for the alternative hypothesis (that is, the variable in question influences the RMT), a Bayes factor > 3 provides moderate and > 10 strong evidence. Conversely, a Bayes factor < 1 provides anecdotal evidence for the null hypothesis (that is, the variable in question does not influence the RMT), a Bayes factor < 0.33 provides moderate and < 0.1 strong evidence (Jeffreys 1961; Lee and Wagenmakers 2014). Bayes factors for a specific fixed effect were assessed by comparing the full model to the model without the factor of interest using the bayestestR package in R (Makowski, Ben-Shachar, and Lüdecke 2019). Bayes factors for correlation coefficients were calculated using the BayesFactor package in R (Morey and Rouder 2015).

## 3 Results

### 3.1 Replication analysis

All study procedures were tolerated well and without side effects. RMT in the dominant hemisphere had a mean of 34.5% (standard deviation 5.9%, range 25-49%). The range of 24% was comparable to Rosso et al. (2017). RMT was positively correlated with CCD (r = 0.626, p < 0.001; Figure 1A). Aligning with Rosso et al. (2017), no correlation was observed between RMT and participants’ age (r = 0.066, p = 0.696; Figure 1B), but the cortical grey matter volume and age (r = −0.557, p < 0.001). However, no meaningful correlation was found between RMT and the cortical grey matter volume of the dominant hemisphere (r = −0.187, p = 0.260; Figure 1C) or FC M1-PMd (r = 0.041, p = 0.805; Figure 1D). There was no association between RMT and FC between any other pair of regions (Table 1).

**Figure 1.**
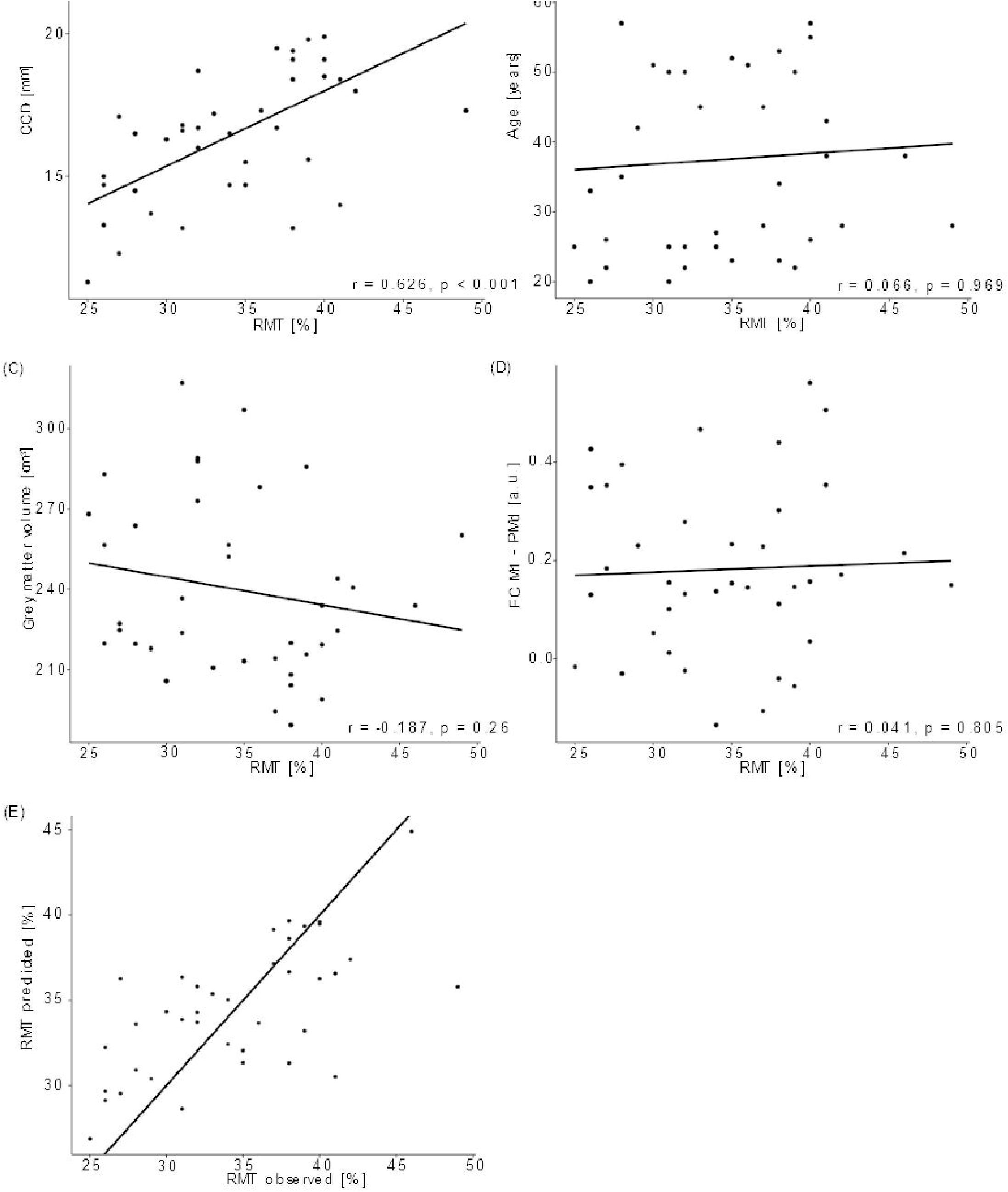
Regression analysis for dominant hemisphere. Correlation between RMT (%) and CCD (A), Age (B), grey matter volume (C) and FC M1-PMd (D). (E) Observed RMT versus RMT predicted by the model. The diagonal line corresponds to perfect prediction.

The multiple regression model explained 42% (R^2^; 95%-CI [23.4%, 65.5%]; Figure 1F) of the variance in RMT. In contrast to Rosso et al. (2017), only CCD was predictive of RMT in this model, while FC M1-PMd and the grey matter volume did not show an effect. Finally, age was not associated with RMT. We obtained strong evidence for the impact of CCD on RMT (BF_10_ = 2.48*10^3^). In contrast, the Bayes factors of the effect of FC M1-PMd (BF_10_ = 0.17), the grey matter volume (BF_10_ = 0.28) and the age (BF_10_ = 0.27) moderately favored the null hypothesis. Detailed results can be found in Table 3.

### 3.2 Extended analysis

The mean RMT for both hemispheres was 34.0% (standard deviation 6.1%, range 23-51%). Comparable to the results for the dominant hemisphere, RMT was positively associated with CCD (estimate: 1.448, p < 0.001; Figure 2A). No association was found with participants’ age (estimate: 0.026, p = 0.708; Figure 2B), cortical grey matter volume (estimate: −0.022, p = 0.445; Figure 2C) and FC M1-PMd (estimate: −0.047, p = 0.986; Figure 2D). Further, the hemisphere stimulated did not impact RMT (estimate: −1.079, p = 0.098; Figure 2E). Again, no association between RMT and FC between any other pair of regions was observed (Table 2).

**Figure 2.**
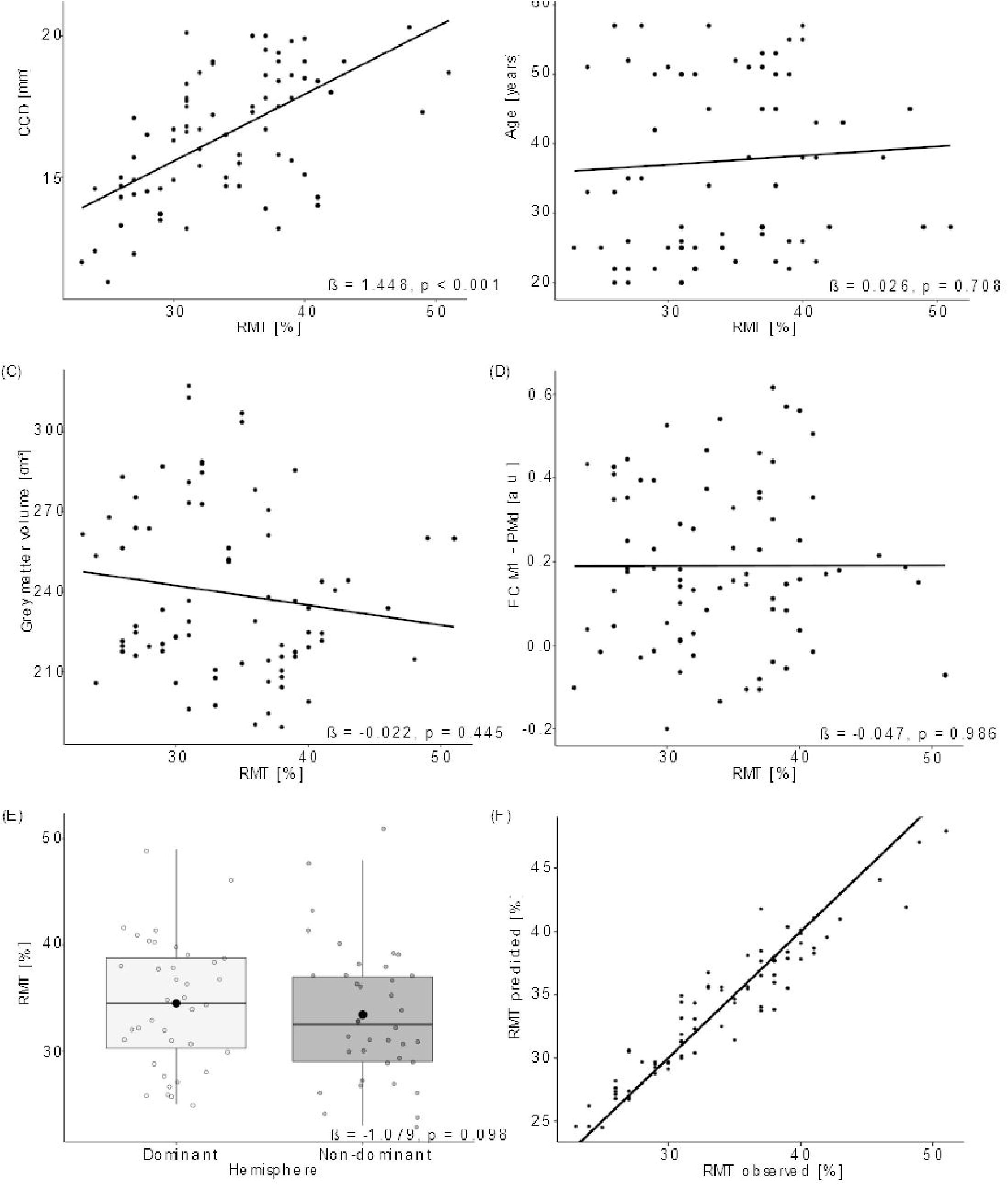
Linear mixed model analysis for both hemispheres. Regression lines between RMT (%) and CCD (A), Age (B), grey matter volume (C) and FC M1-PMd (D). (E) Effect of hemisphere on RMT. Large black dots correspond to the mean RMT for each hemisphere. (F) Observed RMT versus RMT predicted by the model. The diagonal line represents perfect prediction.

The linear mixed model including age, CCD, the cortical grey matter volume and FC M1-PMd explained 44.4% (R^2^; 95%-CI [31.3%, 60.2%]; Figure 2F) of the variance in RMT. Like the multiple regression analysis, CCD was the only significant predictor of RMT. No association was found between RMT and FC M1-PMd, age, the cortical grey matter volume or hemisphere. There was strong evidence for the effect of CCD on RMT (BF_10_ = 1.8*10^4^). In contrast, there was moderate evidence for the null hypothesis when looking at FC M1-PMd (BF_10_ = 0.12), age (BF_10_ = 0.2) and cortical grey matter volume (BF_10_ = 0.16) and anecdotal evidence for the null hypothesis when looking at hemisphere (BF_10_ = 0.42). Detailed results can be found in Table 4.

### 3.3 Analysis of subgroup with successive MRI and TMS

Finally, we repeated these analyses in the subgroup of participants that received their MRI directly before the TMS. The mean RMT for the dominant hemisphere in this subset was 33.1% (standard deviation 5.4%, range 26-39%). The multiple regression model for the dominant hemisphere explained 91% (R^2^; 95%-CI [71.2%, 99.8%]) of the variance in RMT. None of the tested parameters reached significance for predicting RMT (Table 5), which can most likely be explained by the small sample size. We still obtained strong evidence for the impact of CCD (BF_10_ = 93.37) and age (BF_10_ = 142.68) on RMT. In contrast, the Bayes factors of the effect of FC M1-PMd (BF_10_ = 0.39), the grey matter volume (BF_10_ = 0.70) gave anecdotal evidence for the null hypothesis. Importantly, the relationship between RMT and FC M1-PMd estimated here was also positive and thus in the opposite direction compared to Rosso et al. (2017).

**Table 5.** Multiple regression model for the subgroup and dominant hemisphere.

| Dependent variable | Independent variable | Estimate<sup>a</sup> | T value | P value | Partial R<sup>2</sup><sub>a</sub> | BF<sub>10</sub> |
| --- | --- | --- | --- | --- | --- | --- |
| RMT | CCD | 1.661<br>[-0.903, 4.226] | 2.787 | 0.108 | 0.795<br>[0.236, 0.994] | 93.37 |
|  | Age | -0.303<br>[-0.741, 0.134] | -2.983 | 0.096 | 0.816<br>[0.304, 0.995] | 142.68 |
|  | Grey matter volume | -0.033<br>[-0.263, 0.196] | -0.623 | 0.597 | 0.163<br>[0.000, 0.964] | 0.70 |
|  | FC M1-PMd | 1.078<br>[-36.188, 38.345] | 0.124 | 0.912 | 0.008<br>[0.000, 0.951] | 0.39 |

<sup>a</sup>Presented with 95% confidence intervals.

The mean RMT for both hemispheres in this subset was 33.1% (standard deviation 5.2%, range 26-41%). The linear mixed model including data from both hemispheres explained 84.4% (R^2^; 95%-CI [70.1%, 95.1%]) of the variance in RMT. CCD and age were significant predictors of RMT. No association was found between RMT and FC M1-PMd, the cortical grey matter volume or hemisphere. There was strong evidence for the effect of CCD (BF_10_ = 62.07) and age (BF_10_ = 193.89) on RMT and anecdotal evidence for the cortical grey matter volume (BF_10_ = 1.13). In contrast, there was moderate evidence for the null hypothesis when looking at FC M1-PMd (BF_10_ = 0.28) and the hemisphere (BF_10_ = 0.30). Again, the estimated relationship between RMT and FC M1-PMd was positive and thus in the opposite direction compared to Rosso et al. (2017). Detailed results can be found in Table 6.

**Table 6.**
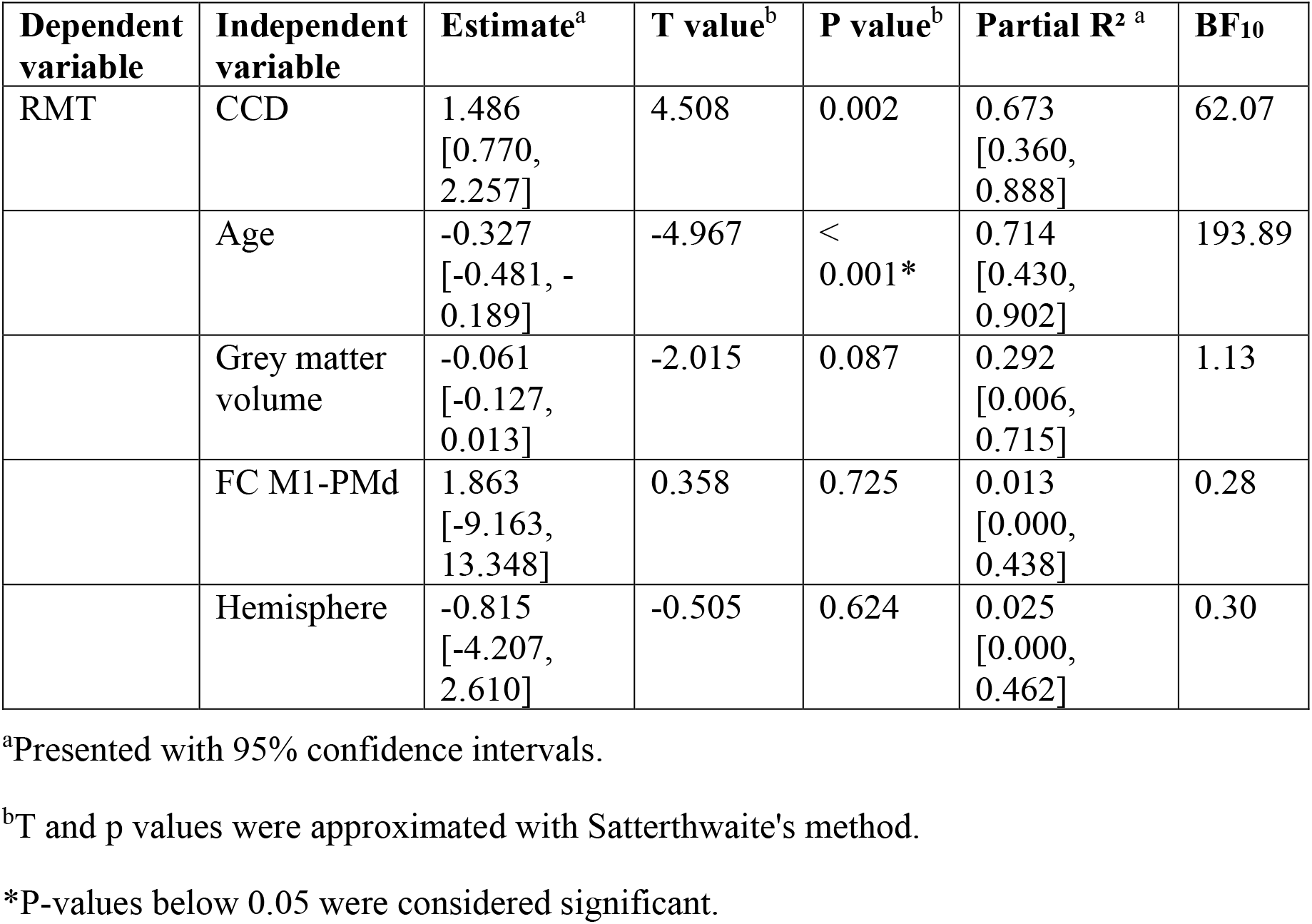
Combined linear mixed model for the subgroup and both hemispheres.

| **Dependent variable** | **Independent variable** | **Estimate<sup>a</sup>** | **T value<sup>b</sup>** | **P value<sup>b</sup>** | **Partial R<sup>2</sup> <sup>a</sup>** | **BF<sub>10</sub>** |
| --- | --- | --- | --- | --- | --- | --- |
| RMT | CCD | 1.486<br>[0.770, 2.257] | 4.508 | 0.002 | 0.673<br>[0.360, 0.888] | 62.07 |
|  | Age | -0.327<br>[-0.481, -0.189] | -4.967 | < 0.001* | 0.714<br>[0.430, 0.902] | 193.89 |
|  | Grey matter volume | -0.061<br>[-0.127, 0.013] | -2.015 | 0.087 | 0.292<br>[0.006, 0.715] | 1.13 |
|  | FC M1-PMd | 1.863<br>[-9.163, 13.348] | 0.358 | 0.725 | 0.013<br>[0.000, 0.438] | 0.28 |
|  | Hemisphere | -0.815<br>[-4.207, 2.610] | -0.505 | 0.624 | 0.025<br>[0.000, 0.462] | 0.30 |

<sup>a</sup>Presented with 95% confidence intervals.
<sup>b</sup>T and p values were approximated with Satterthwaite's method.
\*P-values below 0.05 were considered significant.

## 4 Discussion

The present study aimed to replicate findings by Rosso et al. (2017) on the impact of rsfMRI functional connectivity on RMT. Specifically, Rosso et al. (2017) proposed an influence of FC between M1 and PMd of the dominant hemisphere, while accounting for known predictors such as CCD, cortical grey matter volume and age. In contrast to Rosso et al. (2017), we did not observe an influence of FC between any of the investigated motor regions on RMT in either the dominant hemisphere or when taking into account data from both hemispheres. The absence of this effect was supported by Bayes factors providing moderate evidence for the null hypothesis. The only significant predictor of RMT was CCD, while age, the cortical grey matter volume and the hemisphere had no shown impact on RMT. Notably, our models only explained a maximum of 44% of variance compared to 75% in the study by Rosso et al. (2017) using the same predictors.

The positive association between CCD and RMT due to the exponential decrease of the magnetic field with increasing distance from the coil is well established (McConnell et al. 2001; Kozel et al. 2000; Stokes et al. 2005). Consequently, any factor contributing to an increased distance, such as anatomical variability or brain atrophy, reduces the magnetic field reaching the cortical target areas. To elicit motor evoked potentials comparable in size, the stimulation intensity needs to be increased, leading to a higher RMT in these subjects (McConnell et al. 2001). It has therefore been suggested to measure the RMT in units of the electric field induced at the cortical level rather than percentage of the stimulator output as this should be less susceptible to the cofounding impact of CCD (Julkunen et al. 2012).

Contrary to our expectations, we were not able to observe an effect of age on RMT in the present sample. Others found an increased RMT with age, with aging related brain atrophy, leading to a larger CCD, being the main hypothesized underlying cause (Bhandari et al. 2016; Rosso et al. 2017). However, other studies have – similarly to our findings – reported the absence of an age effect in their samples (Kozel et al. 2000; Wassermann 2002). Similar to age, the cortical grey matter volume was also not predictive of the RMT in our sample. Yet, age and cortical grey matter volume were negatively associated, hinting to the presence of age-related brain atrophy also in our sample.

Rosso et al. (2017) were the first to report an effect of FC between M1 and PMd on RMT. They explained this effect by the known connectivity between both regions and potential facilitatory processes upon stimulation. The present study does not support these conclusions. However, this does not necessarily mean that FC does not impact RMT at all, but rather that such an effect could not be captured using the present methodology. Recent studies (Desideri et al. 2019; Schaworonkow et al. 2019; Zrenner et al. 2018) have shown the state-dependency of TMS-induced effects by investigating the size of motor evoked potentials during different phases of the mu-rhythm observed in human electroencephalography. They showed that stimuli applied to the negative peak of the oscillation cause larger motor evoked potentials compared to the positive peak, thus describing a state of high or low excitability respectively. While functional connectivity using rsfMRI can only be captured at timescales of several seconds (Babiloni et al. 2009; Yaesoubi, Miller, and Calhoun 2017), a similar state-dependency phenomenon might theoretically be observable using this measure. In support of this idea, Tagliazucchi et al. (2012) have related fluctuating FC with spectral power of different oscillation frequencies in electroencephalography, thus underpinning the neurophysiological origin of FC states. Neither the original study (Rosso et al. 2017) nor this replication attempt would have been able to address this state-dependency hypothesis as MRI and TMS were not performed at the same time.

In support of our results, the present study was conducted in a sample almost twice as large as that of Rosso et al. (2017), with additional data from the non-dominant hemisphere. The sample was comparable in terms of participants’ age and gender distribution as well as the range of recorded RMTs. We replicated the statistical analyses of Rosso et al. (2017), while including Bayes factors as a measure to quantify evidence for the respective hypothesis. This is crucial for the current study as it enables us to make assumptions about the null hypothesis (Dienes 2014; 2011; Jeffreys 1961; Lee and Wagenmakers 2014), thus giving evidence for the absence of an effect of FC on RMT. All together, we followed the original protocol as closely as possible with some minor deviations, whose potential impacts on our results will be discussed in the following section.

i. Differences in equipment. Both studies were conducted using a 3T MRI scanner (Siemens AG, Erlangen, Germany) with a 32-channel head coil with almost identical scanning sequences. The rsfMRI sequence in the present study had a slightly shorter TR and larger number of volumes. Similarly, TMS systems differed between both studies (NBS 5, Nexstim: maximal output 1.42 Tesla; Magstim 200^2^, Magstim: maximal output 2.2 Tesla). However, both systems used a neuronavigation software to keep the coil positioning stable and determined RMT manually (Rossini-Rothwell method; Rossini, Barker, and Berardelli 1994; Rothwell et al. 1999). While this impacted the absolute values of RMT (13.5% higher average RMT in the original study compared to this study), the range of RMTs relative to the absolute RMTs was comparable in both studies.
ii. Timing of MRI and TMS. In the study by Rosso et al. (2017), participants received their TMS measurement directly after the MRI scan. In contrast, in the present study the time between both measurements varied, with only seven subjects receiving them directly after another. To address this difference, we included an exploratory analysis for the subgroup of subjects that received the MRI directly before the TMS. It should be noted that this analysis can only give a rough estimate of any potential effect due to the small sample size in this subgroup. There was also no effect of FC on RMT in this analysis. Most rsfMRI networks are fairly reproducible over time (Chou et al. 2012), thus reducing the impact of the time interval between both measurements. On the other hand, varying FC states can be observed even during the short scanning period (Allen et al. 2014; Battaglia et al. 2020; Hutchison et al. 2013; Preti, Bolton, and Van De Ville 2017) and this is further altered by execution of a task such as subject’s movement from MRI to TMS (Gonzalez-Castillo and Bandettini 2018). Thus, also on a theoretical level these factors again seem unlikely to explain deviating results.
iii. Delineation of ROIs. Rosso et al. (2017) used subject-specific ROIs drawn on subjects’ FA maps, while the present study used an atlas. Both approaches lead to comparable ROIs in terms of size and location, with the exception of an additional ROI for the ventral premotor cortex in the atlas used in this study (Mayka et al. 2006). Further, Marrelec and Fransson (2011) show that mean FC values are not impacted by the choice of the ROI delineation method, specifically when resulting differences in ROIs are small.

In conclusion, the present study does not support the concept of functional connectivity between M1 and PMd influencing excitability of the corticospinal tract. The distance between coil and cortex remains the most important factor in explaining variability in RMTs, while other factors like age, grey matter volume or hemisphere seem to be less important. Consequently, results of the present study contradict the hypothesis of RMT reflecting variability of both anatomical and functional features of the motor system as proposed by Rosso et al. (2017). Growing evidence (McConnell et al. 2001; Kozel et al. 2000) highlights the impact of coil to cortex distance and potential impact of other anatomical factors such as microstructural properties of the corticospinal tract (Klöppel et al. 2008). In contrast, more research is needed to investigate the role of functional factors like state-dependency of excitability, wakefulness or the influence of medication. While anatomical factors should remain stable within the same individual over a short period of time and are thus more likely to explain interindividual differences in RMTs, functional factors might be a promising target to explain intraindividual variability of RMT measurements.

## 5 Abbreviations

BF: Bayes factor
CCD: Coil-to-cortex distance
CI: confidence interval
FA: fractional anisotropy
FC: functional connectivity
M1: primary motor cortex
PMd: dorsal premotor cortex
PMv: ventral premotor cortex
RMT: resting motor threshold
ROI: region-of-interest
rsfMRI: resting-state functional magnetic resonance imaging
SMA: supplementary motor area
S1: primary somatosensory cortex
TMS: transcranial magnetic stimulation

## 6 Acknowledgments

This research has been published as a preprint (Engelhardt et al. 2020). We would like to thank Andrea Hassenpflug and Yvonne Kamm for technical support during the MRI scans and the Berlin Center for Advanced Neuroimaging for providing the facilities for conducting the MRI measurements. Further, we thank Dr. Ulrike Grittner for statistical counseling.

## Notes

### Competing Interest Statement

The authors have declared no competing interest.

### Summary of Updates

More details on the analysis of functional connectivity have been added in the methods section to make the approach replicable for the readers. The discussion section has been shortened and condensed. Table descriptions were shortened and moved to footnotes whenever applicable. Correlation coefficients and p-values were added to Figure 1, b-estimates and p-values to Figure 2.

